# Human gaze is systematically offset from the center of cone topography

**DOI:** 10.1101/2021.03.19.436115

**Authors:** Jenny L. Reiniger, Niklas Domdei, Frank G. Holz, Wolf M. Harmening

**Affiliations:** Department of Ophthalmology, Rheinische Friedrich-Wilhelms-Universität Bonn, Bonn, Germany

**Author notes:** Corresponding author: Wolf M. Harmening.

## Abstract

The small physical depression of the human retina, the fovea, is the retinal locus of prime visual resolution, achieved by a peaking topography of the light sensitive cone photoreceptor outer segments ^1–3^ and a post-receptor wiring scheme preserving high-density sampling ^4,5^. Humans dynamically direct their gaze such that the retinal images of objects of interest fall onto the foveola, the central one-degree diameter of the fovea ^6–8^, but it is yet unclear if a relationship between the individual photoreceptor topography at this location and visual fixation behavior exists ^9,10^. By employing adaptive optics in vivo imaging and micro-stimulation ^11–13^, we created topographical maps of the complete foveolar cone mosaics in both eyes of 20 healthy participants while simultaneously recording the retinal location of a fixated visual object in a psychophysical experiment with cellular resolution. We found that the locus of fixation was systematically shifted away from the topographical centers towards a naso-superior quadrant on the retina, about 5 minutes of arc of visual angle on average, with a mirror symmetrical trend between fellow eyes. In cyclopean view, the topographical centers were superior to the fixated target, corresponding to areas in the visual field usually more distant ^14,15^ and thus containing higher spatial frequencies. Given the large variability in foveal topography between individuals, and the surprising precision with which fixation is repeatedly directed to just a small bouquet of cones in the foveola, these findings demonstrate a finely tuned, functionally relevant link between the development of the cellular mosaic of photoreceptors and visual behavior.

## Results and Discussion

### Foveolar cone topography

By high-resolution adaptive optics scanning laser ophthalmoscopy (AOSLO), cone photoreceptor topography at the very center of the fovea was analyzed in 41 eyes of 21 healthy human participants (twenty binocular, one monocular). In each retinal image, about 6800 to 9100 individual cones were resolved, marked and their location used to compute continuous two-dimensional maps of cone density (see Methods, Figure 1A-F). Peak cone density, PCD, varied widely across participants (range: 10,823 – 18,023, average: 14,067 cones/deg^2^, see also: Table S1), similar to previous studies ^1,3,9,10,16–19^. In alignment with histology ^2^ and two *in vivo* imaging studies ^3,17^, we found a steeper drop in cone density along the vertical compared to the horizontal meridian (Figure 1G and 1H). This anisotropy is also found in the retinas of other mammals ^20,21^, and has been discussed to provide a specialized area for scanning the horizon and achieve greater visual acuity and sensitivity to movement^22^.

**Figure 1.**
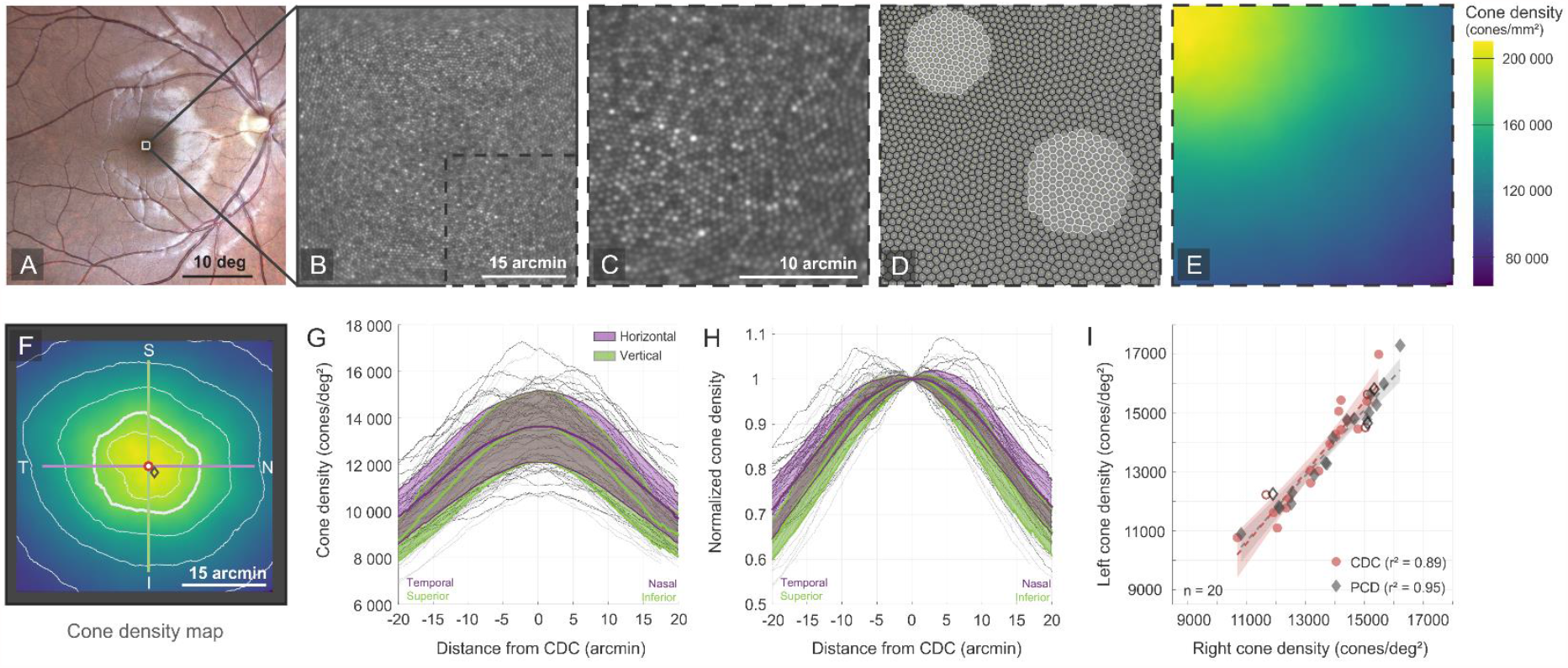
Cellular topography of the foveola. (A) Fundus photograph of a participants’ right eye, with the AOSLO imaging field, 51 arcmin across, shown as white square. (B) Single AOSLO frames were registered and averaged to create fully resolved topographical maps of all foveolar cones (B-F, the dashed quadrant is shown enlarged in C-E). (C) Cone locations were labelled semi-manually by an experienced image grader. (D) Voronoi tesselation allowed for calculation of continuous cone density maps based on the area of the nearest 150 Voronoi patches around each pixel in the image (two examples shown as white cone outlines). (E) Cone density is visualized color coded, expressed here as cones/mm^2^. (F) In the full map, iso-contour lines are 10^th^, 20^th^ (bold), 40^th^, 60^th^ and 80^th^ percentiles. Diamond = peak cone density (PCD), red circle = cone density centroid (CDC). See Figure S1 for maps of all eyes. Green and purple lines indicate the cardinal meridians along which density is shown in G and H for all eyes. (G) Profiles of absolute cone density (40 eyes of 20 participants), visualizing the large range of normal cone densities. (H) The same profiles normalized to the cone density at the CDC. (I) Cone densities at the CDC (red dots) and PCDs (gray diamond) in fellow eyes of 20 participants were highly correlated. Open markers indicate children. Regression lines and 95% confidence intervals are represented by dotted lines and shaded areas, respectively. See Figure S2 for extended symmetry analysis.

During analysis of image material from the same eye across different days, we noticed that the retinal location of peak cone density was quite variable, due to slight variations in local image distortions, image noise and the following cone center annotation. We thus introduce a novel, spatially more robust metric to anchor the topographical center of the fovea: the cone density centroid, CDC, was computed as the weighted center of cone densities within the 20th percentile contour (Figure 1F). In 8 participants (16 eyes), foveolar cone mosaics were imaged and analyzed on two different days. After careful alignment of high signal-to-noise ratio images, the advantage of using the CDC as anchor became apparent. While the PCDs as well as CDC densities were highly correlated between fellow eyes (r^2^ = 0.95, p << 0.001 and r^2^ = 0.89, p << 0.001, respectively), (Figure 1I), on average, PCD locations varied by more than 3-fold (mean ± std: 3.0 ± 2.3 arcmin, range: 0.1 – 7.9 arcmin), compared to CDC locations (mean ± std: 0.9 ± 0.7 arcmin, range: 0.1 – 2.6 arcmin). This difference was statistically significant (paired t-test, p = 0.002). In all following analyses, the CDC was used as the singular spatial reference location of the foveola.

A detailed discussion of factors leading to differences in cone densities reported between studies, e.g. due to the examined population, optical differences in the imaging pathway, imaging wavelength, postprocessing and averaging of the data, different methods that underly cell density calculation, as well as sample preparation (for histological tissue), can be found in the report by Wang et al. (2019) ^3^. The fact that more and more optical and analytical limitations are lifted with novel imaging techniques, like sub diffraction-limited resolution offered by AOSLO ^23^, and the good agreement between *in vivo* studies, is likely to lead towards a replacement of the gold standard for quantitative cone mosaic analysis, from histology of dissected tissue preparations ^2^ towards high resolution *in vivo* imaging.

The three children in our study were aged 10, 12 and 14 years (P3, P10 and P16, respectively) and their PCDs as well as topographical symmetry (see below) did not differ from the adult population. Previous studies found no correlation between age and cone density for participants between 10 and 69 years, ^24^ nor for children between 5.8 and 15.8 years ^25^, at eccentricities between 0.2 mm and 1.5 mm. Our results extend those findings into the foveal center. Histological studies point to an earlier cessation of centripetal cone photoreceptor migration, with a doubling of cone density between gestational week 22 and postnatal day 5 examinations and a tripling between 5 days and 45 months post-natal ^26^. With an average of 108,439 cones/mm^2^, cone density at the age of 45 months equates to about 60% of the average cone density, 183,560 cones/mm^2^, found in our population. Visual acuity in children was shown to approach adult performance between the age of 5 and 6 years ^27^. Thus, the children examined here are assumed to be in a comparable stage of visual development as adults.

### Interocular symmetry of foveolar topography

Symmetry is an extensively studied characteristic in various organs. Previous observations in the field of ophthalmic optics showed ocular symmetries between fellow eyes, such as corneal topography and ocular wavefront aberrations ^28^. For cone density, high interocular correlation was shown at larger retinal eccentricities (250, 420, 760 and 1300 µm) ^29^ as well as for PCD ^17^. In the foveal center, similar as shown in our data, Cava et al. found that, in addition to the PCD, the Voronoi cell area regularity and certain iso-density contour areas are also highly symmetrical between fellow eyes ^1^. Here, we observed high topographical symmetry between fellow eyes, readily perceivable by eye (see Figure S1). When the point-wise difference in density was computed between fellow eyes, the median RMS (3.8 %) was only slightly larger than the difference between two maps of the same eye analyzed from different days (median RMS: 2.9 %) (see supplemental information, Figure S2). Small local image distortions are likely to occur due to the scanning nature of the AOSLO, pixelwise image acquisition and sequential stabilization processes. With a conservative estimation of such local distortions of up to 3 pixels (≙ 0.3 arcmin), they remain relatively small compared to the magnitude of measured offsets between retinal locations of interest. By manually selecting a reference frame with low distortions in our analysis pipeline (see Methods), we further minimized this confound.

Overall, the shape of cone density profiles was similar across subjects (Figure 1G, H) but some eye pairs appeared to have a more particular two-dimensional topography than the average (Figure S1). PCDs in fellow eyes were strongly correlated and not different between right and left eyes (paired t-test, p = 0.6), as is also observed by Cava et al. ^1^. There was also no significant difference of PCDs between dominant and non-dominant eyes in our population (paired t-test, p = 0.4). Preliminary data from an acuity study of our group, including pilot data of five participants from the present study, showed that resolution acuity was better in the dominant eyes of all five examined participants, while acuity thresholds were highly correlated with the density of the foveolar cone mosaic ^30^. This suggests that better performance in the dominant eye might be related to other factors than PCD, e.g. the particular retinal locations used during the task as well as retinal motion. To test this hypothesis, resolution acuity and ocular dominance need to be investigated in a larger population. Additionally, a spatially resolved analysis of retinal image quality might help to better understand how optical limits during development influence the formation of the optimal retinal locus, as they affect the sampling limit in resolution tasks ^31^.

### Preferred retinal locus of fixation

In the natural environment, fixation, discrimination or resolution requirements are often closely related. For a long time, it was common view that the anatomical center of the fovea also represents the center of fixation ^32^, a view supported by the rough alignment between these retinal loci. With current imaging techniques however, opening the door to the exact cellular makeup of an individual eye, it was revealed that the preferred retinal locus of fixation (PRL) is offset from the location of the PCD as well as from the center of the foveal avascular zone and the foveal pit ^3,9,10,16,33^. The PRL is also not the retinal location that provides highest sensitivity to small spot stimuli, which was recently shown to be rather plateau-like within the central 0.1 degree of the foveola ^34^. Together, these findings leave a possible systematic relationship between the PRL and the retinal cone mosaic open.

In the majority of eyes in our population (33/41), fixation behavior was examined on two or more days (see Table S1). PRLs could be found accurately, with a median distance of 2.3 arcmin between consecutive measurements (range: 1.0 - 5.6 arcmin) (Figure 2B and Figure S3B). When stimulus locations were pooled across a single day (≥ 3 × 150 video frames pooled), median locations differed by only 1.5 arcmin (range: 1.0 - 4.2 arcmin) (Figure S3A, B). The observed fixation stability, given by the Isoline areas, ISOA, ranged between 23 and 153 arcmin^2^ in right eyes, and between 29 and 154 arcmin^2^ in left eyes (Table S1). The participants who had a larger median ISOA also had higher PRL variability between single measurements (ρ = 0.39, p = 0.01) (Figure 2C). PRLs as well as ISOAs were highly reproducible in individuals, even across a period of up to 3.5 years (Figure S3A). This confirms and extends the finding of Kilpeläinen et al., showing PRL reproducibility over a period of two days, on average ^33^.

**Figure 2.**
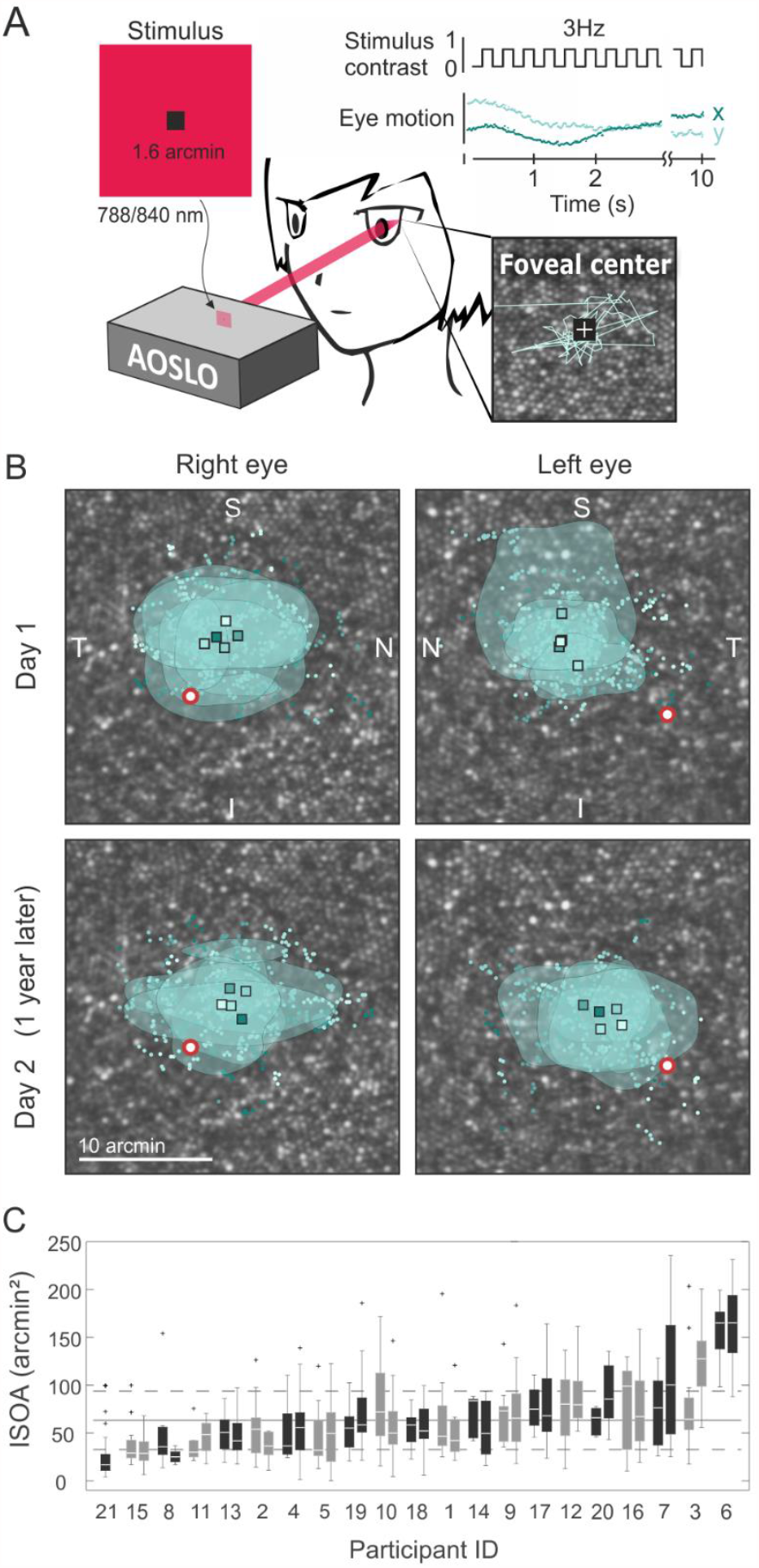
Measuring fixation behavior. (A) Participants fixated a small flashing black square presented in the center of the AOSLO imaging raster (top left). High-resolution eye motion traces were derived from at least 5 consecutive, 10 sec AOSLO videos (top right). (B) The PRLs, computed as two-dimensional median of all stimulus locations, are shown for consecutive measurements in the right eye (left column) and left eye (right column) of P5 on two different days (∼1 year between measurements, see Figure S3 for fixation stability across multiple years) as square marker. Small dots indicate the retinal location of the stimulus in individual video frames, increasing marker brightness represents consecutive videos. Contours are the area containing one standard deviation of the data (Isoline area, ISOA). The CDC is indicated by a red circle with white fill. (C) ISOAs in all eyes per participants, ordered by their average magnitude. The left bars represent left and right bars right eyes, respectively. The solid and dotted lines represent the mean value of all data and one standard deviation, respectively. Box whisker extend to the most extreme data values and plus markers represent outliers (distance from box > 1.5 × range between 25^th^ and 75^th^ percentile).

Small differences in PRL variations between measurement days (Kilpeläinen et al.: 0.52 arcmin; this study: 1.5 arcmin) might be observed because of differences in the fixational tasks. Kilpeläinen et al. used a moving target, possibly providing higher fixation stability due to a higher attentional demand for the participant ^35^. In addition to the structural symmetry between fellow eyes described in the previous section, we also observed functional symmetries. Albeit recorded under monocular viewing, fixation stability across fellow eyes was highly correlated (r^2^ = 0.66, p << 0.001, Figure 2C), supporting the hypothesis of an underlying coupling of both eyes during fixation ^36^. When eyes were grouped according to ocular dominance, there was no difference between median ISOAs of dominant and non-dominant eyes (p = 0.062, Wilcoxon signed rank test, n = 20). Previous studies found functional interocular correlation in microsaccade rates and amplitudes under monocular viewing conditions ^37^, bivariate contour ellipse areas ^38^, and suggest improved fixation stability under binocular viewing conditions ^36,37^.

### The relationship of cone topography and fixation

By measuring fixation behavior in a cone-resolved experiment (Figure 2A and B) and by careful alignment with the cone density maps of both eyes, we here reveal a fine and very reproducible systematic offset between cone topography and fixation behavior. In retinal coordinates, the PRL was displaced naso-superiorly from the CDC by an average amount of 4.7 arcmin (Figure 3A), corresponding to about 10 cone diameters. A recent monocular study reported a comparable distance between PRL and PCD location (average: 5.08 arcmin) ^33^. Other studies, with a lower number of subjects or a less accurate method of measuring the PRL, found larger offsets with median values of 9.8 and 11.5 arcmin ^9,10^. However, a trend towards a PRL superior to the PCD is visible also in those data.

**Figure 3.**
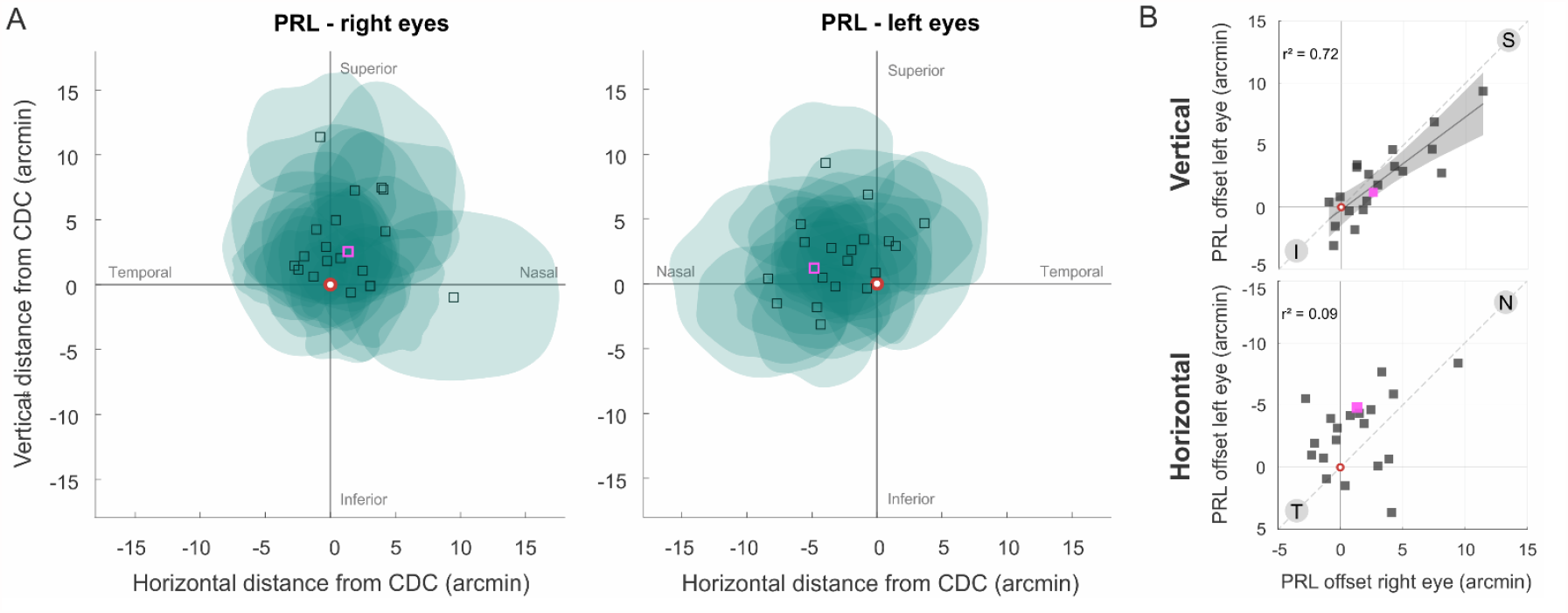
Relationship between cone topography and PRL in retinal coordinates. (A) Right and left eyes’ median PRL and 1 STD ISOAs plotted relative to the CDC (see also Figure S4). As an example, the PRL offsets of the participant (P13) whose offsets were most similar to the average are highlighted in purple. (B) The vertical offset components were strongly correlated between fellow eyes (r^2^ = 0.72, p << 0.001). The regression line and the 95 % confidence interval are represented by a solid line and gray shaded area, respectively. The horizontal component showed a non-significant correlation between fellow eyes (r^2^ = 0.09, p = 0.19). The dotted lines represent a perfectly mirror symmetrical offset between both eyes. S=superior, I=inferior, N=nasal, T=temporal retinal orientations. See Figure S4 for distance and angular relationships.

The median PRLs observed in the present study omit the temporal-inferior area, even though some of the ISOAs reach into this quadrant as well. In one participant (P19) only, both PRLs were offset temporally (Figure 3B and Figure S1). The average angular distance between CDC and PRL was 4.6 and 5.0 arcmin in right and left eyes, respectively.

Similar PRL offsets (4.7 and 5.9 arcmin) could be found when re-analyzing the data from Wang et al., 2019 ^3^ with our analysis methods (Figure S4A). The offset distances were correlated between fellow eyes in our study (r^2^ = 0.45, p = 0.001, Figure S4B), with a high impact resulting from a strong correlation in the vertical direction (r^2^ = 0.72, p << 0.001, Figure 3B). Horizontal offsets were not significantly correlated (r^2^ = 0.09, p = 0.19), thus, the offset’s angular component was not significantly correlated (r^2^ = 0.07, p = 0.28), albeit with a mirror symmetrical trend (Figure S4C). The data provided by Wang et al. showed similar but not significant trends, which might be due to a lower accuracy in PRL measurements (at least one 10 sec video), as we show that pooling data over multiple consecutive videos reveals more reproducible PRL locations (Figure S3B).

### Projection of retinal correspondence to the visual field

In the following, we assume that PRLs are identical under monocular and binocular viewing conditions and that PRLs are retinal coordinates of corresponding points in the visual environment and therefore fixed positions independent of the viewing distance ^39^. In the cyclopean view, with the PRLs of both eyes as common center, the CDCs were slightly superior to the fixated point (Figure 4A). Given the interindividual variations of cone topography and the here evaluated size of high-cone-density areas (see Methods), this yielded an overall orbital high-density distribution that was, on average, shifted to being centered slightly above the fixated target (Figure 4B). In a natural environment, the visual field above a fixated point is often farther away as there are many horizontal surfaces (e.g. grounds and table tops) ^14,15^. Most surfaces are textured, creating a bottom-to-top gradient of spatial frequencies with higher frequencies above and lower frequencies below the point of fixation. Even though the superior displacement of the CDC was relatively small, it might allow for a better estimation of the 3D structure of such textured surfaces or other objects in the natural environment. Due to the strong correlation of the vertical position in the visual field, the offset between both eyes CDCs was more pronounced horizontally in our data (Figure 4C). In the cyclopean view, the CDCs of most right eyes landed leftward from the left eye’s CDCs, resulting in crossed disparity. Uncrossed disparities were observed in five participants only, who also had minimal horizontal offsets. Figure 4D further illustrates the geometry and magnitude of the small offsets projected into the binocular field of view. We note that our analysis is based on observations during prolonged fixation maintenance, a laboratory condition rather unusual in natural viewing ^40^, since resolution is enhanced by retinal motion ^41–43^. Reading, for instance, is a highly dynamic process with ongoing crossing and uncrossing of the line of sights. Binocular coordination during reading was found to be biased towards more uncrossed disparities by about 1 character ^44^. In an attempt to extrapolate our offsets to such highly dynamic situations, this could imply that for the majority of fixations during reading, the centers of cone topography move closer together than for the case of maintained fixation, thereby enlarging the binocularly overlapping area of high cone densities. This might be beneficial for reading tasks, as it was also shown that reading is more difficult with monocular than with binocular viewing ^45,46^.

**Figure 4.**
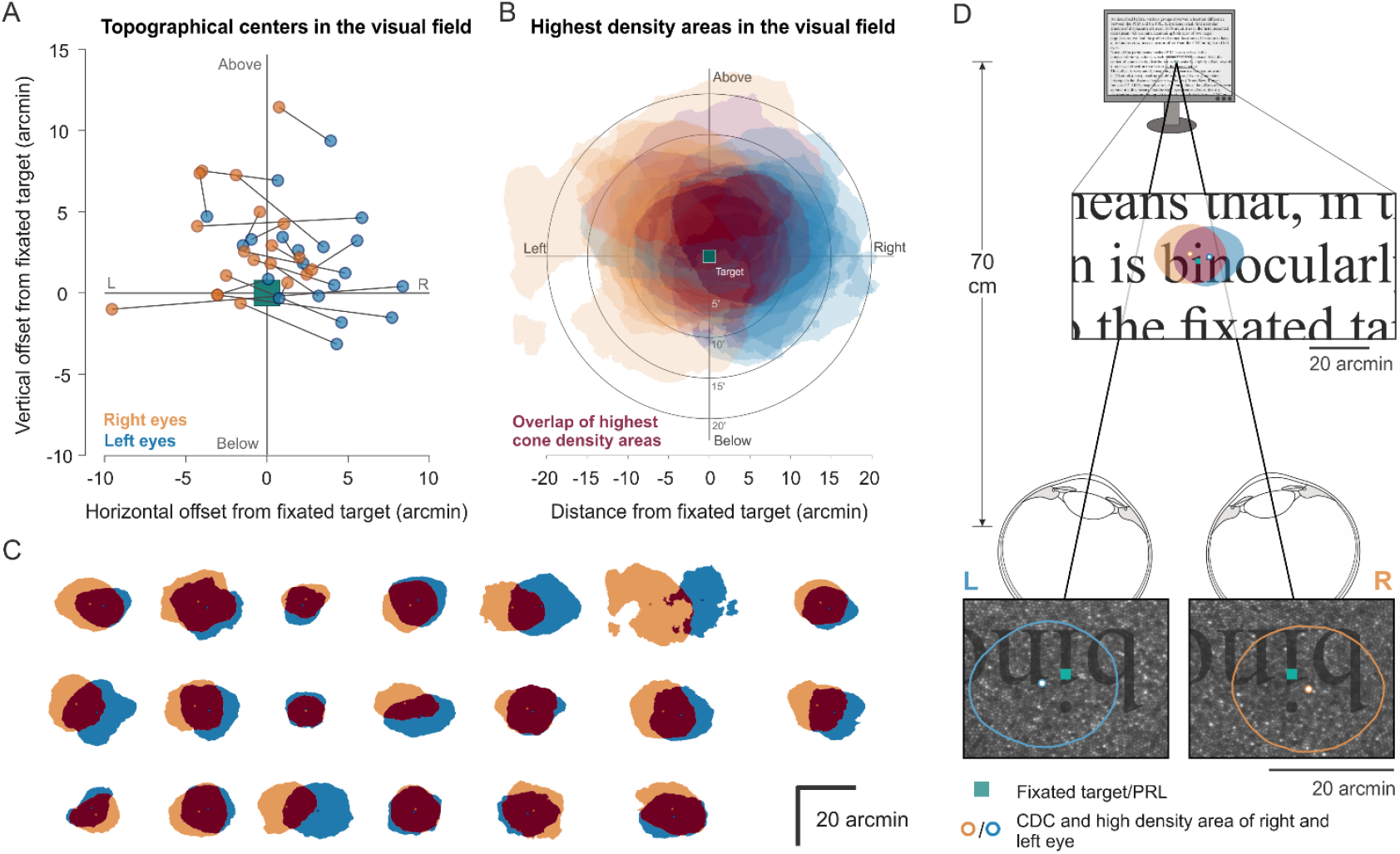
Projection of cone topography into the binocular visual field. (A) The CDC of both eyes (right = orange, left = blue) plotted with their PRL as common center in the visual field. The lines connect fellow eyes. (B) Similar as in A, including the retinal areas encompassing the highest 20 % of cone densities (within the central 50 arcmin of the fovea). Overlap is shown in dark red. High density areas are, on average, slightly offset towards details superior to the fixated object and displaced horizontally between eyes. (C) Projections of high-density cone areas of all participants. (D) The average horizontal displacement between the two eyes, in a static reading situation, roughly equals the distance between two letters at reading distance (Times New Roman, fontsize 12, 100% magnification, ≙ 1 mm). The orientation of cone mosaics is as seen from behind the observer.

The highly ordered and systematic functional and topographical architecture we observe between fellow eyes could be the result of a developmental process creating appropriate location information for binocular spatial sampling. From such point of view, a nasal displacement might emerge as an “overshoot” during PRL formation, ensuring overlap between the high spatial frequency sampling capacity areas in both eyes. We found incomplete overlaps which essentially enlarge the visual field sampled by high-density cones in all participants (Figure 4B, C). By the rules of binocular combination, the eye that sees higher contrast and sharper details gets more weight in the cyclopean percept ^47^. Thus, by imperfect horizontal alignment of cone topographies, the visual system might create a larger field of sharp perception with individual sharpness gradients of the two retinal images ^48^. One of the factors driving the enrichment of visual capacities during development is the demand of resolving fine structures in the visual environment. Apart from photoreceptor sampling, the topography of retinal connectivity within the foveolar midget circuit might also be important: the connectivity between individual cones and midget bipolar and ganglion cells was recently shown to develop and establish a private line for the central photoreceptors already during gestation ^4^. The centripetal migration of cone photoreceptors starts in parallel, but takes place mainly after birth ^26^. The nasal superior offset direction aligns with the closest connectivity to the optic nerve head, which could facilitate the slightly offset PRL development, even if conduction velocity of retinal ganglion cells was shown to minimize possible time differences across the retina ^49^. At larger retinal eccentricities, midget ganglion cells have smaller dendritic field diameters in the nasal quadrant of human retinae ^50^, which may be an outcome of the same underlying mechanisms as the biased PRL formation.

## Conclusions

Taken together, participants without known retinal disease or abnormalities showed a small but systematic offset between their PRL and the center of cone density distribution, formed in a way to vertically offset high cone densities towards the superior part of the visual field and to ensure a horizontal overlap of those areas in the binocular visual field. This functional symmetry was associated with high interocular symmetry of foveolar cone topography. Binocular, foveated display systems that seek to mimic human vision with high precision could be tuned to reflect this spatial relationship ^51^. Binocular *in vivo* foveal topography data may provide a basis for detecting changes in the central photoreceptor topography during retinal disease ^52^, and more generally, could contribute to replace histology as the gold standard for normative human photoreceptor evaluations in a healthy population.

## Acknowledgements

We thank Austin Roorda and the members of his lab for kindly providing their image and PRL data for re-analysis. We are grateful for insightful comments by the reviewers of an earlier version of this manuscript.

This work was supported by the Carl Zeiss Foundation (HC-AOSLO), and the Emmy Noether Program of the German Research Foundation (DFG, Ha 5323/5-1).

## Author contributions

J.L.R and W.M.H conceived the research idea. J.L.R, N.D and W.M.H developed the data analysis pipeline. J.L.R performed the data analysis and convolutional neural network training. J.L.R and W.M.H wrote the manuscript. All authors discussed the results and edited the manuscript.

## Methods

### Participant Details

Forty-one eyes of twenty-one participants (7 male, 14 female, 38 adults [age: 18 – 42], 3 children [age: 10, 12 and 14]) with no known ocular conditions and only mild refractive errors (SE: ± 2.5 diopters) were studied. Participants are referred to throughout the manuscript with a singular ID, selected based on a descending order of peak cone densities for the left eye. For one of the participants (P21), data from the right eye were included as the only monocular dataset in the study, because the left eye’s cone mosaic could not be resolved completely. Therefore, this eye’s data were only used for PRL reproducibility analysis, as image and functional data were collected over multiple years. Most of the participants were examined on multiple days (compare Table S1). Participants P4, P13 and P21 were trained AOSLO observers and members of the lab. Mydriasis was established by two drops of 1% Tropicamide, instilled into the eyelid about 15 and 10 minutes prior to the imaging session. A third drop was administered in case imaging and experimentation continued for more than 30 minutes. A customized dental impression mold (bite bar) was used to immobilize and adjust the head position and thus to align the participants eye in front of the imaging system. Written informed consent was obtained from each participant and all experimental procedures adhered to the tenets of the Declaration of Helsinki, in accordance with the guidelines of the independent ethics committee of the medical faculty at the Rheinische Friedrich-Wilhelms-Universität of Bonn.

### Adaptive optics retinal imaging

*In vivo* images of the complete foveolar cone mosaic were recorded using a custom-built adaptive optics scanning laser ophthalmoscope (AOSLO). The general setup of the AOSLO has been described previously ^53,54^, pertinent differences are described here. Briefly, the AOSLO front-end featured three f = 500 mm afocal telescopes, designed to point-scan an adaptive optics corrected focal spot of light across the retina to achieve diffraction limited resolution performance in both the incident and reflected beams. A magnetic actuator-driven deformable mirror with continuous membrane surface (DM97-07, 7.2 mm pupil diameter, ALPAO, Montbonnot-Saint-Martin, France) was placed in a retinal conjugate plane and driven by the error signals of a 25×25 lenslet Shack Hartmann sensor (SHSCam AR-S-150-GE, Optocraft GmbH, Erlangen, Germany). Imaging and wavefront correction wavelength was either 840 nm (± 12 nm) or 788 nm (± 12 nm) light, obtained by serial dichroic and bandpass filtering of a supercontinuum source (SuperK Extreme EXR-15, NKT Photonics, Birkerød, Denmark). The imaging field of view was 0.85 × 0.85 degree of visual angle. The light reflected from the retina was captured in a photomultiplier tube (PMT, H7422-50, Hamamatsu Photonics, Hamamatsu, Japan), placed behind a confocal pinhole (Pinhole diameter = 20 µm, equaling 0.47 (840nm) and 0.5 (788nm) Airy disk diameters). The PMT signal was sampled continuously in a field programmable gate array (FPGA), rendering a 512 × 512 pixel video at 30 Hz (600 pixel per degree of visual angle). With fast acousto-optic modulation of the imaging wavelengths, the square imaging field becomes a retinal display in which psychophysical visual stimulation was possible ^11,55^. The best images during PRL recordings (see below) were used to create spatially registered, high signal to noise ratio images of the foveal center in which all cones could be resolved.

### Image processing and cone density analysis

Acquired AOSLO video frames were spatially stabilized by real-time, strip-wise image registration in custom written software ^56^. These online-stabilized videos contained frames displaying incomplete stabilization that could be due to poor image quality, eye blinks, drying tear film, etc. Such frames were identified and deleted manually. The remaining frames were averaged to obtain a single high-quality image of each retina per video. The single best of at least five such images was selected to be used for further analysis and serve as high signal-to-noise anchor image for spatial alignment with functional data recordings. All cone center locations were labeled in a semi-manual process by a single trained image grader: first, a convolutional neural network ^57^, CNN, was trained to locate cone center locations with a smaller subset of only manually graded images in our pilot study. Then, all retinal images were annotated by the newly trained CNN, and manually corrected using custom software. Such corrections were especially necessary in the foveal center, and wherever cones appeared completely dark ^58^. The manual correction prioritized mosaic regularity in cases of ambiguity ^2^. Based on the labeled cone center locations, a Voronoi tessellation was computed (MATLAB functions: *delaunayTriangulation, voronoiDiagram* and *voronoin*). Each cone was regarded as occupying the space of each corresponding Voronoi cell. Angular cone density (cones/deg^2^) was computed at each image pixel by averaging the Voronoi area of the nearest 150 encircled cones around that pixel (Figure 1D). This method ensured smooth cone density maps and prevented sampling artifacts as they often occur using defined shapes of masks (e.g. circular or square masks) for selection of cones in a particular area (Figure 1E). Linear cone densities were computed with respect to the individual retinal magnification factors of each eye, considering axial length, anterior chamber depth and corneal curvature ^16^, based on swept source biometry (IOLMaster 700, Carl Zeiss Meditech, Jena, Germany). Finally, the cone density centroid (CDC) was determined as the weighted centroid (MATLAB function: *regionprops(region_logical, image, “WeightedCentroid”)*) of the highest 20 % of cone density values. The CDC is indicated by circular marker throughout the manuscript. The 20^th^ percentile was chosen arbitrarily because the entire contour was evaluable in all eyes. CDC locations did only marginally change at other contours. At the 20% contour, cone densities equaled the highest ∼13 % of densities across the entire retina, considering previously reported cone densities at larger retinal eccentricities ^2,19^. Therefore, the theoretical limit of cone sampling within those areas was < 35 arcsec (range: 9600 to 14900 cones/deg^2^), equaling 20/13 vision or better under correction of ocular aberrations. Under natural viewing conditions, expected performance would be slightly less due to higher order aberrations and crucially depends on post-receptor circuitry, midget bipolar and ganglion cells (see Discussion).

To quantify overall symmetry between density maps, three different analyses were performed:

1. Spatial two-dimensional differences (or reproducibility) of cone density maps of the same eye were recorded and analyzed independently on different days (columns in Figure S2A and 2B), based on a careful alignment of the cone mosaic images.
2. The differences between density maps of fellow eyes which were recorded on the same day (rows in Figure S2A and 2B) were obtained by comparing flipping the left eyes map along the vertical axis and aligning it with the CDC of the right eyes map.
3. The difference between individual density maps of all right eyes and a randomly selected left eye, which was flipped and aligned with the CDC of the right eye as described in analysis (2).

To quantify the two-dimensional differences, the root-mean-square (RMS) of the point-by-point difference maps was used for the comparison between absolute and normalized density maps (Figure S2C, respectively, compare with Figure 1G and 1H).

### Determination of the preferred retinal location of fixation (PRL)

Using the AOSLO as stimulation platform, a small (nominal 1.6 arcmin), flashing (3 Hz) square with negative contrast polarity (light turned off) was presented as visual target at the center of the AOSLO imaging raster during image acquisition, and participants were asked to fixate the target as accurately and relaxed as possible. At least five 10-second AOSLO videos were recorded in each eye during such fixation epochs. In AOSLO videos, the visual stimulus was directly visible with respect to the retina (Figure 2A and 3B). Thus, fixation behavior can be directly and unambiguously observed in such videos. The PRL was calculated as the median fixation target location across all videos. To bring fixation behavior into spatial correspondence with the topographical analysis, averaged retinal images derived from both analyses independently were carefully aligned with each other. In 33 of the 41 eyes, PRL measurements were conducted multiple times (e.g. if participants also took part in other experiments). In three eyes (P13, P4 and P21), data were obtained in 8, 12 and 17 measurement sessions, respectively, over a period of 3.5 years. For eight participants (16 eyes) sessions were repeated after 1 year. For quantification of fixation stability, the isoline area (ISOA) which contains one standard deviation (STD) of the data was fitted to the scatterplot of all stimulus positions (Figure S2B).

## Supplemental Information

**Figure S1.**
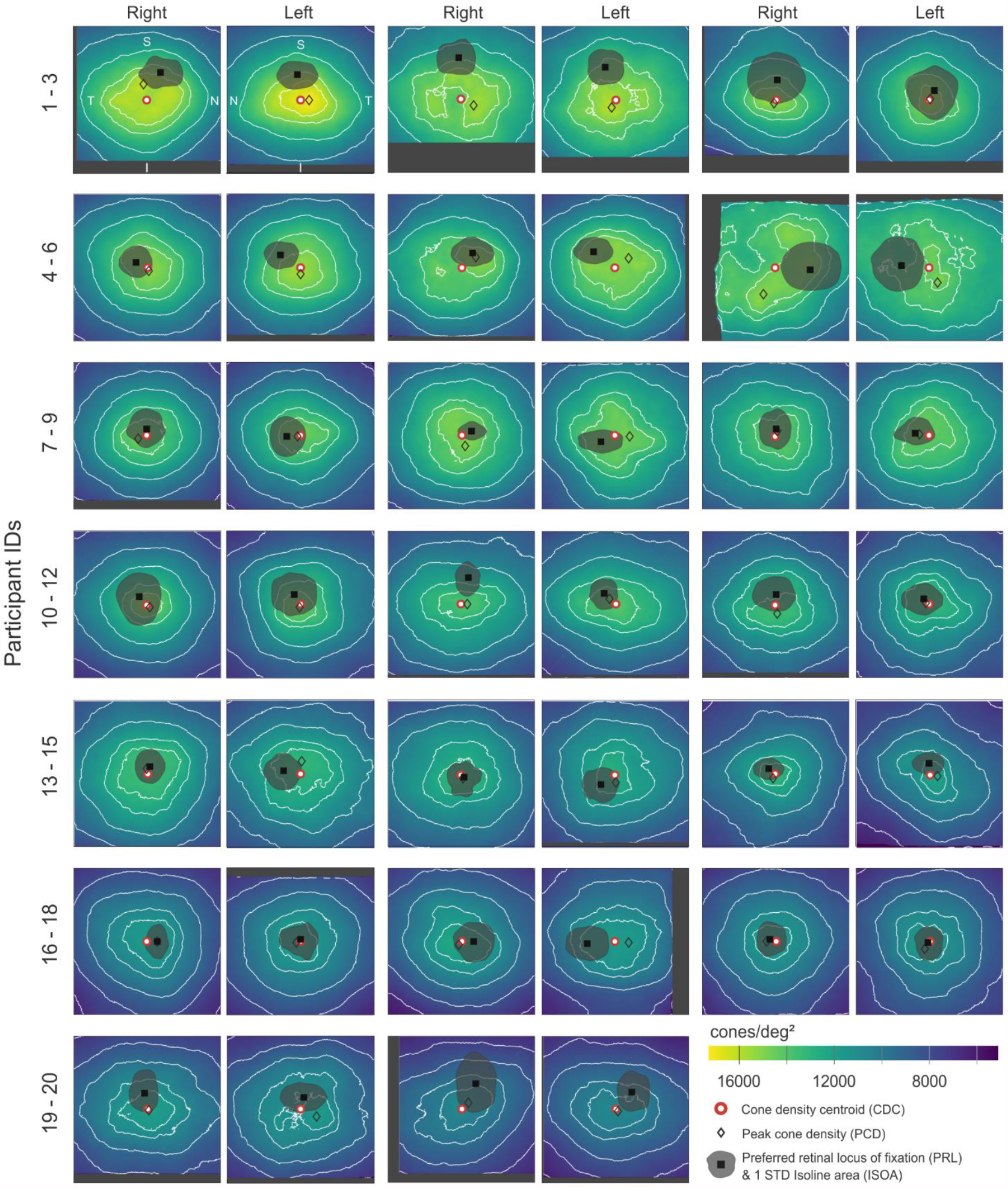
Binocular cone density contour maps of all participants. The central 40 × 40 arcmin density maps of right and left eyes are presented in fundus orientation for all participants (Superior retina is up, nasal is right for right eyes, and left in left eyes). Participants IDs (P1-P20) were ordered by PCD value exhibited, from top left to bottom right in this representation. Iso-contour lines represent the 10^th^, 20^th^, 40^th^, 60^th^ and 80^th^ percentile of density values. The cone density centroid (CDC) is indicated by a red circle. PCD = diamond, PRL = square, shown in the center of the individual fixation isoline areas (1 STD). Dark gray streaks reflect parts of the image that were cropped because of borders or poor quality.

**Figure S2.**
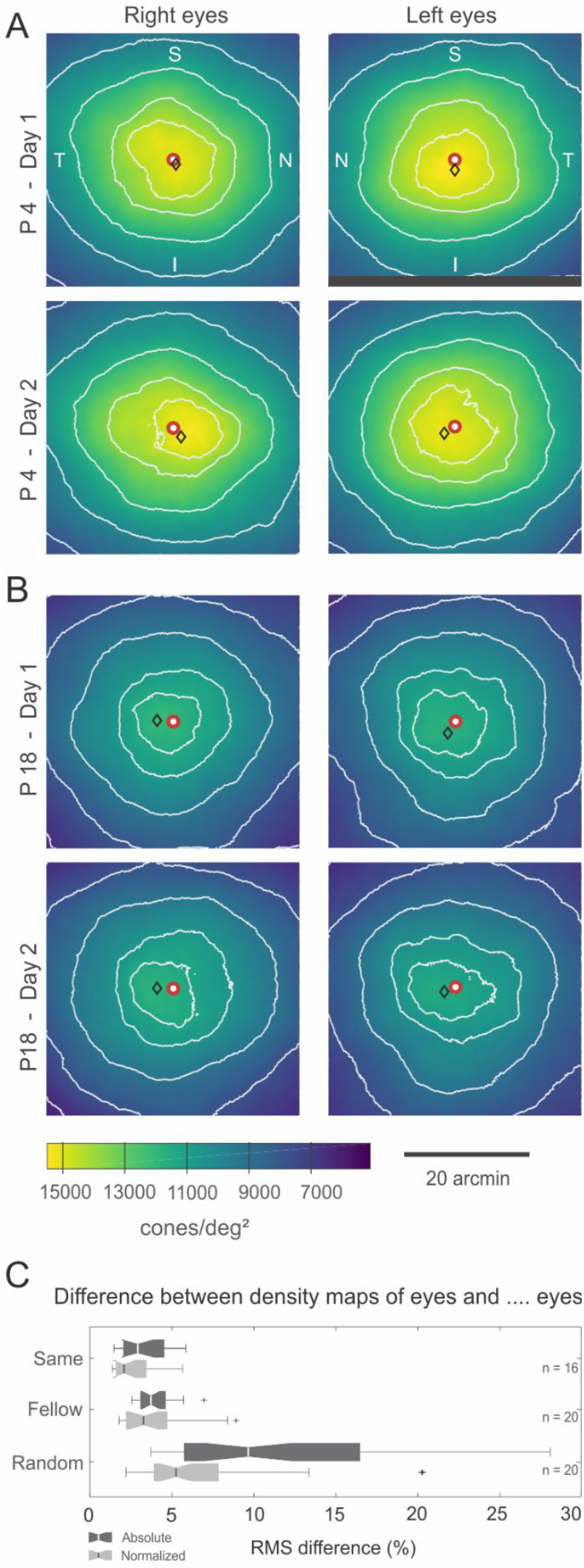
Symmetry of foveolar cone density maps. (A+B) Cone density maps of both eyes of two participants (P4 and P18) are shown in fundus orientation. Rows represent imaging sessions on different days. Contour line definition and marker (CDC and PCD) as in Figure 1. (C) The root mean square (RMS) of differences in absolute cone density values (dark fill) was evaluated for: (1) same eyes on 2 different days (median: 2.9 %, range: 1.5 – 5.9 %), (2) fellow eyes on the same day (median: 3.8 %, range: 2.6 – 6.9 %) and (3) the comparison between individual eyes and random fellow eyes showed the greatest differences (median: 9.6 %, range: 3.7 – 28.1 %), that can be explained to a large extend by the variation in absolute density values among participants. To focus more on the two-dimensional shape of the maps, the normalized density maps were compared in the same way: (1) reproducibility showed a median value of 2.1 % (range: 1.4 – 5.6 %), (2) symmetry differences between fellow eyes were slightly greater (median: 3.3 %, range: 1.8 – 8.9 %) and the normalized differences between random partner eyes were highest (median: 5.3 %, range: 2.2 – 13.4 %). The notch represents the 95% confidence interval of the median, box whiskers extend to the most extreme data values and plus markers represent outliers (distance from box > 1.5 × range between 25th and 75th percentile).

**Figure S3.**
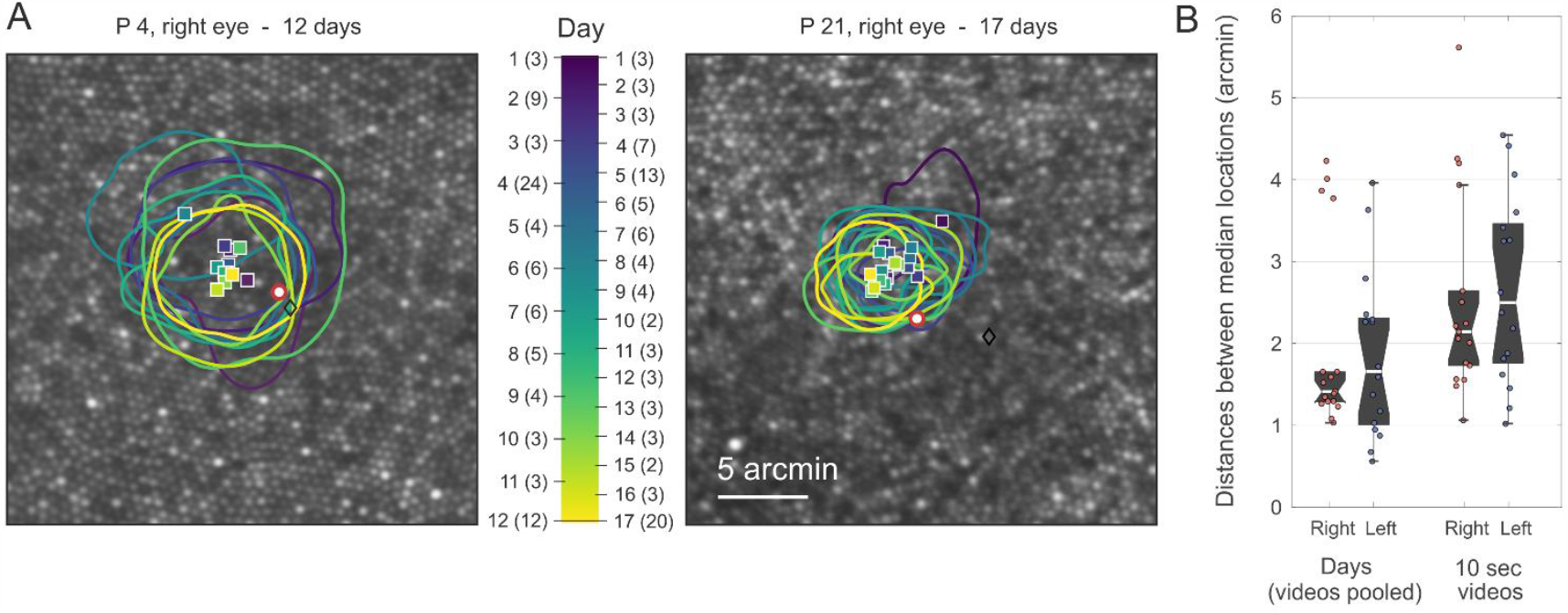
Fixation stability across multiple years. (A) PRL measurements in two participants (P4 and P21) were recorded over a period of 3.5 years on 12 and 17 different days, respectively. Color represents measurement sessions and thus time between first and last examination (2017-2020). Small dots are individual stimulus locations, squares are PRLs shown inside their 1 STD isoline areas. (B) The effect of data pooling compared between data taken from individual 10 sec videos and when pooled across multiple of such 10 sec videos recorded at the same day in all studied eyes.

**Figure S4.**
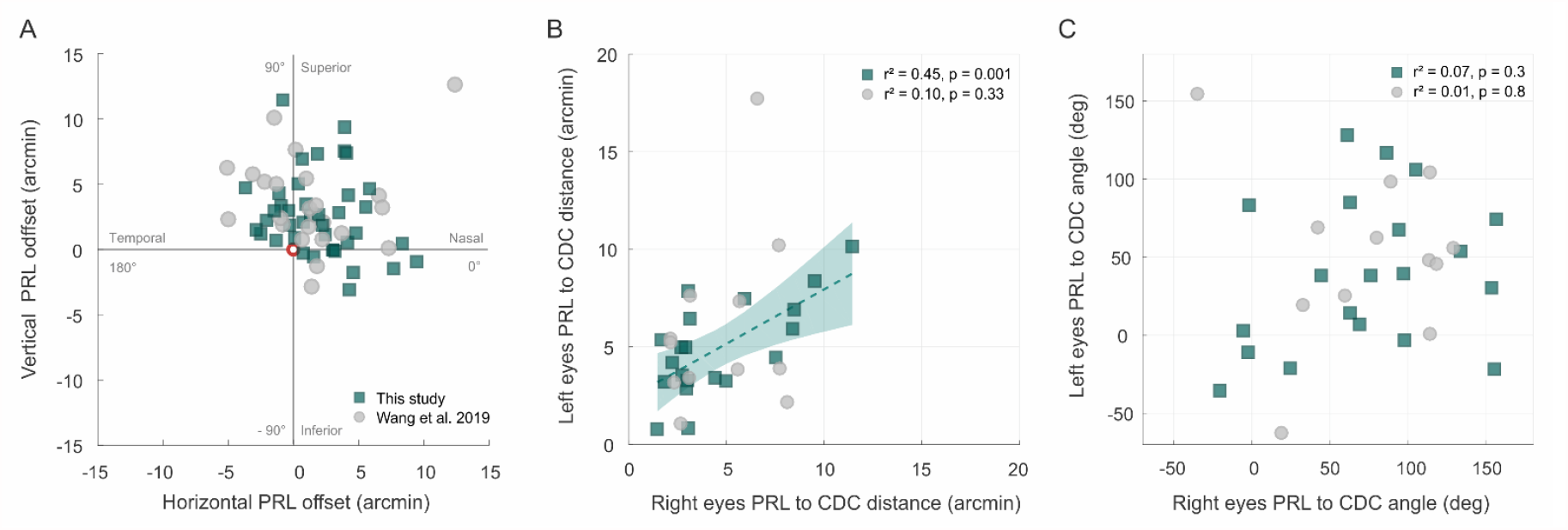
PRL offsets in this study and re-analyzed data from Wang et al. (2019). (A) The right and left eyes’ PRL relative to the CDC combined for both eyes (this study = squares, Wang et al. = circles). (B) Offset distance was significantly correlated between fellow eyes in our study (r^2^ = 0.41, p = 0.002), but not in the data provided by Wang et al. (r^2^ = 0.1, p = 0.33). (C) While demonstrating a similar trend, offset direction did not show a significant correlation between fellow eyes in either population (this study: r^2^ = 0.05, p = 0.36; Wang et al.: r^2^ = 0.01, p = 0.76).

**Table S1.** Participant statistics, ocular dominance, foveal cone mosaic metrics and PRL details. Ocular dominance was measured prior to PRL measurements. PCD and CDC densities are given in angular and linear units (retinal magnification was computed based on axial length, anterior chamber depth and retinal curvature for each eye ^16^). The PRL was measured on different days and PRL variability is the retinal distance between single repeated PRL measurements (runs), or pooled data across days. Note that in participant 21, the complete cone mosaic could only be resolved in the right eye.

| Participant ID | Gender | Age | Eye | Dominance | Cone mosaic |  |  |  | Fixation behavior |  |  |  |  |  |
| --- | --- | --- | --- | --- | --- | --- | --- | --- | --- | --- | --- | --- | --- | --- |
|  |  |  |  |  | PCD<br>cones/deg<sup>2</sup> | PCD<br>cones/mm<sup>2</sup> | Density<br>@CDC<br>cones/deg<sup>2</sup> | Density<br>@CDC<br>cones/mm<sup>2</sup> | Total # of<br>videos | #<br>Measure-<br>ment<br>days | PRL<br>variability<br>across<br>single runs<br>(arcmin) | PRL<br>variability<br>across<br>days<br>(arcmin) | Median<br>ISOA<br>(arcmin<sup>2</sup>) | Median<br>ISOA<br>across<br>days<br>(arcmin<sup>2</sup>) |
| 1 | M | 22 | L | 0 | 17309 | 210145 | 16997 | 206354 | 8 | 2 | 3.6 | 3.6 | 70.8 | 68.8 |
|  |  |  | R | 1 | 16217 | 196477 | 15488 | 187643 | 8 | 2 | 5.6 | 3.9 | 50.3 | 128.8 |
| 2 | F | 21 | L | 0 | 16003 | 214584 | 15454 | 207225 | 10 | 2 | 3.3 | 4.0 | 54.9 | 45.7 |
|  |  |  | R | 1 | 15667 | 210212 | 14217 | 190754 | 10 | 2 | 4.2 | 1.3 | 33.8 | 97.9 |
| 3 | M | 10 | L | 1 | 15814 | 217811 | 15650 | 215550 | 12 | 2 | 4.5 | 1.4 | 63.5 | 153.9 |
|  |  |  | R | 0 | 15350 | 212760 | 15124 | 209632 | 12 | 2 | 3.9 | 4.2 | 126.4 | 183.0 |
| 4 | F | 29 | L | 0 | 15657 | 193441 | 15429 | 190622 | 15 | 3 | 1.9 | 2.3 | 36.6 | 46.9 |
|  |  |  | R | 1 | 15230 | 190708 | 15101 | 189087 | 82 | 12 | 2.2 | 1.2 | 52.4 | 56.7 |
| 5 | M | 30 | L | 1 | 15301 | 207076 | 15058 | 203793 | 15 | 3 | 1.8 | 2.4 | 32.0 | 46.8 |
|  |  |  | R | 0 | 15428 | 205507 | 14137 | 188313 | 15 | 3 | 1.5 | 1.6 | 66.1 | 61.0 |
| 6 | F | 28 | L | 1 | 15018 | 172925 | 14282 | 164448 | 10 | 2 | 3.3 | 1.6 | 154.4 | 157.3 |
|  |  |  | R | 0 | 15157 | 170318 | 14030 | 157655 | 11 | 2 | 2.6 | 1.1 | 152.6 | 166.8 |
| 7 | F | 23 | L | 1 | 14828 | 198349 | 14774 | 197625 | 15 | 3 | 2.6 | 0.9 | 88.0 | 101.0 |
|  |  |  | R | 0 | 14662 | 196147 | 14453 | 193346 | 15 | 3 | 2.2 | 1.3 | 93.0 | 114.0 |
| 8 | F | 30 | L | 0 | 14792 | 190964 | 14474 | 186860 | 10 | 2 | 2.4 | 2.3 | 38.1 | 53.8 |
|  |  |  | R | 1 | 15102 | 188851 | 14789 | 184940 | 10 | 2 | 1.8 | 1.3 | 25.1 | 27.2 |
| 9 | F | 26 | L | 1 | 14763 | 207763 | 14433 | 203125 | 10 | 2 | 1.5 | 0.6 | 73.2 | 61.2 |
|  |  |  | R | 0 | 14426 | 205894 | 14215 | 202877 | 9 | 2 | 2.5 | 1.4 | 66.9 | 86.6 |
| 10 | M | 12 | L | 0 | 14657 | 201831 | 14514 | 199867 | 13 | 2 | 3.4 | 1.2 | 70.9 | 107.5 |
|  |  |  | R | 1 | 15126 | 207682 | 15035 | 206433 | 10 | 2 | 4.2 | 3.8 | 47.0 | 112.3 |
| 11 | F | 26 | L | 1 | 14175 | 187061 | 13971 | 184364 | 5 | 1 | 1.6 | - | 30.0 | 49.9 |
|  |  |  | R | 0 | 13938 | 182418 | 13836 | 181085 | 6 | 1 | 2.1 | - | 48.3 | 47.5 |
| 12 | F | 26 | L | 0 | 13315 | 150790 | 13069 | 148002 | 10 | 2 | 1.0 | 1.0 | 64.3 | 70.0 |
|  |  |  | R | 1 | 13697 | 153715 | 13191 | 148042 | 10 | 2 | 2.2 | 1.0 | 76.4 | 86.0 |
| 13 | M | 32 | L | 0 | 13286 | 145870 | 13046 | 143232 | 16 | 3 | 4.1 | 2.8 | 44.7 | 53.6 |
|  |  |  | R | 1 | 13733 | 148503 | 13470 | 145658 | 37 | 8 | 2.0 | 1.6 | 40.7 | 49.1 |
| 14 | F | 20 | L | 0 | 12897 | 176732 | 12640 | 173207 | 5 | 1 | 1.1 | - | 83.8 | 72.2 |
|  |  |  | R | 1 | 13302 | 183331 | 13175 | 181574 | 5 | 1 | 2.8 | - | 49.5 | 64.1 |
| 15 | F | 29 | L | 0 | 12304 | 158973 | 12108 | 156444 | 15 | 3 | 1.6 | 0.7 | 29.0 | 39.4 |
|  |  |  | R | 1 | 13381 | 169702 | 12449 | 157885 | 24 | 5 | 1.1 | 1.4 | 23.3 | 32.2 |
| 16 | M | 14 | L | 0 | 12259 | 162728 | 12233 | 162376 | 5 | 1 | 2.8 | - | 98.9 | 71.6 |
|  |  |  | R | 1 | 11904 | 158460 | 11682 | 155512 | 5 | 1 | 2.4 | - | 67.2 | 43.5 |
| 17 | F | 23 | L | 1 | 11927 | 171912 | 11772 | 169675 | 15 | 3 | 2.2 | 2.3 | 75.0 | 92.4 |
|  |  |  | R | 0 | 12524 | 179022 | 12363 | 176715 | 36 | 6 | 2.3 | 1.6 | 65.3 | 85.5 |
| 18 | F | 31 | L | 0 | 11818 | 163501 | 11617 | 160723 | 15 | 3 | 1.2 | 1.0 | 62.1 | 53.3 |
|  |  |  | R | 1 | 12105 | 165061 | 11939 | 162803 | 15 | 3 | 1.6 | 1.3 | 55.9 | 57.6 |
| 19 | F | 24 | L | 0 | 11789 | 161489 | 11089 | 151900 | 10 | 2 | 4.4 | 1.7 | 53.3 | 73.2 |
|  |  |  | R | 1 | 12131 | 168226 | 12053 | 167150 | 10 | 2 | 2.1 | 4.0 | 62.5 | 64.5 |
| 20 | F | 18 | L | 0 | 10894 | 159556 | 10761 | 157603 | 5 | 1 | 3.2 | - | 66.0 | 78.9 |
|  |  |  | R | 1 | 10823 | 157559 | 10692 | 155656 | 5 | 1 | 4.4 | - | 85.4 | 141.9 |
| 21 | M | 42 | R | 1 | 18023 | 221889 | 16601 | 204383 | 87 | 17 | 1.6 | 1.5 | 16.7 | 22.8 |
| Median |  |  |  |  | 14426 | 183331 | 14030 | 181574 | 10 | 2 | 2.3 | 1.5 | 62.5 | 68.8 |

## References

1. Cava, J.A., Allphin, M.T., Mastey, R.R., Gaffney, M., Linderman, R.E., Cooper, R.F., and Carroll, J. (2020). Assessing Interocular Symmetry of the Foveal Cone Mosaic. Investig. Opthalmology Vis. Sci. 61, 1–11.

2. Curcio, C.A., Sloan, K.R., Kalina, R.E., and Hendrickson, A.E. (1990). Human photoreceptor topography. J. Comp. Neurol. 292, 497–523.

3. Wang, Y., Bensaid, N., Tiruveedhula, P., Ma, J., Ravikumar, S., and Roorda, A. (2019). Human foveal cone photoreceptor topography and its dependence on eye length. Elife 8, 1–21.

4. Zhang, C., Kim, Y.J., Silverstein, A.R., Hoshino, A., Reh, T.A., Dacey, D.M., and Wong, R.O. (2020). Circuit Reorganization Shapes the Developing Human Foveal Midget Connectome toward Single-Cone Resolution. Neuron 108, 905–918.

5. Bryman, G.S., Liu, A., and Do, M.T.H. (2020). Optimized Signal Flow through Photoreceptors Supports the High-Acuity Vision of Primates. Neuron 108, 335–348.

6. Poletti, M., Listorti, C., and Rucci, M. (2013). Microscopic eye movements compensate for nonhomogeneous vision within the fovea. Curr. Biol. 23, 1691–1695.

7. Poletti, M., Rucci, M., and Carrasco, M. (2017). Selective attention within the foveola. Nat. Neurosci. 20, 1413–1417.

8. Ko, H.K., Poletti, M., and Rucci, M. (2010). Microsaccades precisely relocate gaze in a high visual acuity task. Nat. Neurosci. 13, 1549–1554.

9. Putnam, N.M., Hofer, H.J., Doble, N., Chen, L., Carroll, J., and Williams, D.R. (2005). The locus of fixation and the foveal cone mosaic. J. Vis. 5, 632–639.

10. Wilk, M.A., Dubis, A.M., Cooper, R.F., Summerfelt, P., Dubra, A., and Carroll, J. (2017). Assessing the spatial relationship between fixation and foveal specializations. Vision Res. 132, 53–61.

11. Harmening, W.M., Tuten, W.S., Roorda, A., and Sincich, L.C. (2014). Mapping the perceptual grain of the human retina. J. Neurosci. 34, 5667–5677.

12. Rossi, E.A., and Roorda, A. (2010). The relationship between visual resolution and cone spacing in the human fovea. Nat Neurosci 13, 156–157.

13. Sincich, L.C., Zhang, Y., Tiruveedhula, P., Horton, J.C., and Roorda, A. (2009). Resolving single cone inputs to visual receptive fields. Nat. Neurosci. 12, 967–969.

14. Sprague, W.W., Cooper, E.A., Tošić, I., and Banks, M.S. (2015). Stereopsis is adaptive for the natural environment. Sci. Adv. 1.

15. Gibaldi, A., and Banks, M.S. (2019). Binocular eye movements are adapted to the natural environment. J. Neurosci. 39, 2877–2888.

16. Li, K.Y., Tiruveedhula, P., and Roorda, A. (2010). Intersubject variability of foveal cone photoreceptor density in relation to eye length. Invest. Ophthalmol. Vis. Sci. 51, 6858–6867.

17. Zhang, T., Godara, P., Blanco, E.R., Griffin, R.L., Wang, X., Curcio, C.A., and Zhang, Y. (2015). Variability in Human Cone Topography Assessed by Adaptive Optics Scanning Laser Ophthalmoscopy. Am. J. Ophthalmol. 160, 290–300.

18. Cooper, R.F., Wilk, M.A., Tarima, S., and Carroll, J. (2016). Evaluating Descriptive Metrics of the Human Cone Mosaic. Investig. Opthalmology Vis. Sci. 57, 2992–3001.

19. Wells-Gray, E.M., Choi, S.S., Bries, A., and Doble, N. (2016). Variation in rod and cone density from the fovea to the mid-periphery in healthy human retinas using adaptive optics scanning laser ophthalmoscopy. Eye 30, 1135–1143.

20. Packer, O., Hendrickson, A.E., and Curcio, C.A. (1989). Photoreceptor topography of the retina in the adult pigtail macaque (Macaca nemestrina). J. Comp. Neurol. 288, 165–183.

21. Wikler, K.C., Williams, R.W., and Rakic, P. (1990). Photoreceptor mosaic: Number and distribution of rods and cones in the rhesus monkey retina. J. Comp. Neurol. 297, 499–508.

22. Rowe, M.H., and Stone, J. (1976). Properties of ganglion cells in the visual streak of the cat’s retina. J. Comp. Neurol. 169, 99–125.

23. Lu, R., Aguilera, N., Liu, T., Liu, J., Giannini, J.P., Li, J., Bower, A.J., Dubra, A., and Tam, J. (2021). In vivo sub-diffraction adaptive optics imaging of photoreceptors in the human eye with annular pupil illumination and sub-Airy detection. Optica 8, 333–343.

24. Park, S.P., Chung, J.K., Greenstein, V., Tsang, S.H., and Chang, S. (2013). A study of factors affecting the human cone photoreceptor density measured by adaptive optics scanning laser ophthalmoscope. Exp. Eye Res. 108, 1–9.

25. Mirhajianmoghadam, H., Jnawali, A., Musial, G., Queener, H.M., Patel, N.B., Ostrin, L.A., and Porter, J. (2020). In vivo assessment of foveal geometry and cone photoreceptor density and spacing in children. Sci. Rep. 10, 1–14.

26. Yuodelis, C., and Hendrickson, A. (1986). A qualitative and quantitative analysis of the human fovea during development. Vision Res. 26, 847–855.

27. Lai, Y.H., Wang, H.Z., and Hsu, H.T. (2011). Development of visual acuity in preschool children as measured with Landolt C and Tumbling e charts. J. AAPOS 15, 251–255.

28. Francisco Castejón-Mochón, J., López-Gil, N., Benito, A., and Artal, P. (2002). Ocular wave-front aberration statistics in a normal young population. Vision Res. 42, 1611–1617.

29. Lombardo, M., Lombardo, G., Schiano Lomoriello, D., Ducoli, P., Stirpe, M., and Serrao, S. (2013). Interocular symmetry of parafoveal photoreceptor cone density distribution. Retina 33, 1640–1649.

30. Reiniger, J.L., Domdei, N., Linden, M., Holz, F.G., and Harmening, W.M. (2019). Relationship between the foveal photoreceptor mosaic and adaptive optics corrected visual acuity. In Investigative Opthalmology & Visual Science (ARVO Annual Meeting Abstract), p. 1777.

31. Reiniger, J.L., Lobecke, A.C., Sabesan, R., Bach, M., Verbakel, F., de Brabander, J., Holz, F.G., Berendschot, T.T.J.M., and Harmening, W.M. (2019). Habitual higher order aberrations affect Landolt but not Vernier acuity. J. Vis. 19, 1–15.

32. Polyak, S. (1949). Retinal structure and colour vision. Doc. Ophthalmol. 3, 24–56.

33. Kilpeläinen, M., Putnam, N.M., Ratnam, K., and Roorda, A. (2020). The Anatomical, Functional and Perceived Location of the Fovea in the Human Visual System. SSRN Electron. J., doi:10.2139/ssrn.3699785.

34. Domdei, N., Reiniger, J.L., Holz, F.G., and Harmening, W.M. (2021). Retinal factors of visual sensitivity in the human fovea. bioRxiv, doi:10.1101/2021.03.15.435507.

35. Bowers, N.R., Gautier, J., Lin, S., and Roorda, A. (2021). Fixational eye movements depend on task and target. bioRxiv, doi: 10.1101/2021.04.14.439841.

36. Krauskopf, J., Cornsweet, T.N., and Riggs, L.A. (1960). Analysis of eye movements during monocular and binocular fixation. J. Opt. Soc. Am. 50, 572–578.

37. González, E.G., Wong, A.M.F., Niechwiej-Szwedo, E., Tarita-Nistor, L., and Steinbach, M.J. (2012). Eye position stability in amblyopia and in normal binocular vision. Investig. Ophthalmol. Vis. Sci. 53, 5386–5394.

38. Zhu, X., He, W., Du, Y., Zhang, K., and Lu, Y. (2019). Interocular symmetry of fixation, optic disc, and corneal astigmatism in bilateral high myopia: The Shanghai high Myopia study. Transl. Vis. Sci. Technol. 8, 1–11.

39. Hillis, J.M., and Banks, M.S. (2001). Are corresponding points fixed? Vision Res. 41, 2457– 2473.

40. Poletti, M., and Rucci, M. (2016). A compact field guide to the study of microsaccades: Challenges and functions. Vision Res. 118, 83–97.

41. Rucci, M., Iovin, R., Poletti, M., and Santini, F. (2007). Miniature eye movements enhance fine spatial detail. Nature 447, 851–854.

42. Ratnam, K., Domdei, N., Harmening, W.M., and Roorda, A. (2017). Benefits of retinal image motion at the limits of spatial vision. J. Vis. 17, 1–11.

43. Rucci, M., Ahissar, E., and Burr, D. (2018). Temporal Coding of Visual Space. Trends Cogn. Sci. 22, 883–895.

44. Liversedge, S.P., White, S.J., Findlay, J.M., and Rayner, K. (2006). Binocular coordination of eye movements during reading. Vision Res. 46, 2363–2374.

45. Heller, D., and Radach, R. (1999). Eye Movements in Reading: Are two eyes better than one? In Current Oculomotor Research: Physiological and psychological aspects, W. Becker, H. Deubel, and T. Mergner, eds. (New York: Plenum Press), pp. 341–348.

46. Kirkby, J.A., Webster, L.A.D., Blythe, H.I., and Liversedge, S.P. (2008). Binocular Coordination During Reading and Non-Reading Tasks. Psychol. Bull. 134, 742–763.

47. Home, R. (1978). Binocular summation: A study of contrast sensitivity, visual acuity and recognition. Vision Res. 18, 579–585.

48. Gibaldi, A., Labhishetty, V., Thibos, L.N., and Banks, M.S. (2021). The blur horopter: Retinal conjugate surface in binocular viewing. J. Vis. 21, 1–21.

49. Stanford, L.R. (1987). Conduction velocity variations minimize conduction time differences among retinal ganglion cell axons. Science 238, 358–360.

50. Dacey, D.M. (1993). The mosaic of midget ganglion cells in the human retina. J. Neurosci. 13, 5334–5355.

51. Tan, G., Lee, Y.-H., Zhan, T., Yang, J., Liu, S., Zhao, D., and Wu, S.-T. (2018). Foveated imaging for near-eye displays. Opt. Express 26, 25076–25085.

52. Song, H., Rossi, E.A., and Williams, D.R. (2021). Reduced foveal cone density in early idiopathic macular telangiectasia. BMJ Open Ophthalmol. 6, 1–4.

53. Roorda, A., Romero-borja, F., Iii, W.D., and Qneener, H. (2002). Adaptive optics scanning laser ophthalmoscopy. Opt. Express vol, 10 pp 405–412.

54. Domdei, N., Domdei, L., Reiniger, J.L., Linden, M., Holz, F.G., Roorda, A., and Harmening, W.M. (2018). Ultra-high contrast retinal display system for single photoreceptor psychophysics. Biomed. Opt. Express 9, 157–172.

55. Poonja, S., Patel, S., Henry, L., and Roorda, A. (2005). Dynamic visual stimulus presentation in an adaptive optics scanning laser ophthalmoscope. J Refract Surg 21, 575–580.

56. Arathorn, D.W., Yang, Q., Vogel, C.R., Zhang, Y., Tiruveedhula, P., and Roorda, A. (2007). Retinally stabilized cone-targeted stimulus delivery. Opt. Express 15, 13731–13744.

57. Cunefare, D., Fang, L., Cooper, R.F., Dubra, A., Carroll, J., and Farsiu, S. (2017). Open source software for automatic detection of cone photoreceptors in adaptive optics ophthalmoscopy using convolutional neural networks. Sci. Rep. 7, 1–11.

58. Bruce, K.S., Harmening, W.M., Langston, B.R., Tuten, W.S., Roorda, A., and Sincich, L.C. (2015). Normal Perceptual Sensitivity Arising From Weakly Reflective Cone Photoreceptors. Invest. Ophthalmol. Vis. Sci. 56, 4431–4438.

